# Lysosomal function, resistance to stress and repair are compromised by expression of the Alexander disease GFAP R239C mutant

**DOI:** 10.1101/2024.12.03.626547

**Authors:** Elena Hernández-Gerez, Nuria Goya-Iglesias, Álvaro Viedma-Poyatos, María A. Pajares, Dolores Pérez-Sala

## Abstract

Intermediate filaments are critical regulators of cell responses and organizers of cellular structures. GFAP (glial fibrillary acidic protein) is an intermediate filament protein mainly expressed in astrocytes. GFAP mutations are associated with Alexander disease (AxD), a type of leukodystrophy that causes degeneration of astrocytes and ultimately neurodegeneration. AxD astrocytes display protein aggregation, proteostasis defects and altered organelle homeostasis. We previously showed that expression of GFAP AxD mutants in astrocytes provoked mitochondrial alterations and oxidative stress. Here we have used an astrocytoma cell model to explore the impact of GFAP AxD mutants on the lysosomal degradation pathway. Expression of GFAP AxD mutants in this model elicits marked alterations in lysosomal distribution. Cells expressing the GFAP R239C mutant display defective lysosomal activity and intraluminal acidification. Lysosomes are primary sites of oxidative damage. Moreover, expression of GFAP R239C increases susceptibility of lysosomes to oxidative stress, resulting in a greater loss of lysosomal “mass” and compromised membrane integrity, as revealed by increased intraluminal recruitment of galectins, with respect to cells expressing GFAP wt. Notably, lysosomes in GFAP R239C expressing cells are also more vulnerable to chemically-induced rupture. Interestingly, lysosomes of cells expressing GFAP wt are able to rapidly recover after removal of the damaging agent. In sharp contrast, recovery of acidic vesicles is severely impaired in cells expressing GFAP R239C, suggesting a defect in lysosomal repair. Taken together, our results show that expression of the GFAP AxD mutant is sufficient to deeply perturb lysosomal distribution, function and repair. These alterations could contribute to proteostasis defects and cellular toxicity in AxD.

## Introduction

Alexander’s disease (AxD) is a type of leukodystrophy, a disorder that results in the degeneration of the white matter and eventually in neurodegeneration. AxD is a rare disease that frequently debuts before 2 years of age (early onset), but that may also debut later in life (late onset). The disease may manifest with megalocephalia, seizures, spasticity, and/or cognitive impairment, and sometimes leads to a fatal outcome [1–3]. This pathology is caused by single point mutations of the glial fibrillary acidic protein (GFAP), a type III intermediate filament protein that is mainly expressed in astrocytes [3]. In healthy astrocytes, GFAP forms a homogeneous network of filaments, which heteropolymerize with vimentin [1, 4], another type III intermediate filament protein, and extend from the nuclear periphery towards the plasma membrane. In contrast, GFAP mutants associated with AxD form abnormal networks characterized by the presence of bundles and are present in complex aggregates of characteristic appearance, known as Rosenthal fibers [5, 6].

GFAP is involved in multiple cell functions, including astrocyte motility, proliferation and interaction with neurons [7]. Moreover, studies in AxD models have shown that expression of GFAP mutants associates with anomalies in organelle distribution, morphology and function [8–10]. Among them, AxD iPSC-derived astrocytes present alterations in the endoplasmic reticulum structure and in the size and distribution of lysosomes [8]. Morphological alterations of mitochondria have also been observed in several experimental systems [9, 10], as well as impairment of mitochondrial transfer between AxD HESC-derived astrocytes, and from these astrocytes to neuronal cells [11]. Interestingly, expression of certain AxD GFAP mutants (e.g. R239C) in an astrocytoma cell line is sufficient to elicit morphological and functional alterations of mitochondria, with increased production of mitochondrial reactive oxygen species, which could contribute to the generation of oxidative stress in those cells [9].

Oxidative stress is a common trait of multiple neurodegenerative diseases [12–15]. Importantly, GFAP is itself a target of oxidative stress [9, 16, 17], and oxidative and lipoxidative modifications profoundly alter GFAP assembly and organization in cells [9, 16]. These drastic effects are mainly mediated by the modification of the single cysteine residue of GFAP, C294 [16]. Notably, this cysteine residue is conserved in all type III intermediate filaments, that besides GFAP, include vimentin, desmin and peripherin, as well as across species [18]. Interestingly, the high reactivity of this residue together with its functional importance in type III intermediate filament assembly have led us to postulate its role as a redox sensor [16, 17, 19–21]. Importantly, most of the mutations leading to AxD involve changes to more nucleophilic amino acids (mostly cysteine or histidine) [3], which imply gaining new putative oxidation and lipoxidation sites. In fact, some of the most severe forms of AxD are found in patients affected by GFAP mutants with an additional cysteine [3], as in GFAP R239C, which is also one of the mutations most frequently reported. This led us to hypothesize that AxD GFAP mutants could be more susceptible to modification and disruption by oxidants and electrophiles. Indeed, we observed GFAP R239C crosslinks dependent on cysteine oxidation in cellular AxD models [9], whereas crosslinks of this and other cystine-generating mutants have been recently observed in transfected rat astrocytes as well as in vitro [22]. GFAP R239C is also more vulnerable to lipoxidation [9], and exposure of cells expressing this mutant to oxidants or electrophilic mediators greatly increases its aggregation [9]. Besides oxidative modifications, the organization of the GFAP network is modulated by other posttranslational modifications, including phosphorylation and ubiquitination [22, 23]. Remarkably, both anomalous phosphorylation and ubiquitination of GFAP have been observed in AxD [22, 23]. These modifications could also contribute to GFAP aggregation.

Occurrence of protein aggregates and alterations in proteostasis are hallmarks of neurodegenerative diseases, including Alzheimer’s, Parkinson’s and Huntington’s disease, which are characterized by the accumulations of β-amyloid peptide, α-synuclein and huntingtin, respectively [24]. These protein accumulations reflect the presence of abnormally folded and/or modified proteins as well as the failure of the cellular proteolytic machinery, mainly the proteasome and lysosomes, to dispose of this material. Both clearance systems are frequently compromised in neurodegeneration [25–28].

Importantly, type III intermediate filaments are required for the recruitment of misfolded ubiquitinated proteins to aggresomes upon proteasome inhibition, as well as for the adequate distribution of lysosomes [29, 30]. These organelles are highly sensitive to oxidative stress. Lysosomes are naturally rich in iron, and certain forms of iron are capable of catalyzing the splitting of hydrogen peroxide giving rise to the highly reactive hydroxyl radical by the Fenton reaction [31]. Therefore, under oxidative stress, high levels of hydrogen peroxide can reach the lysosome and undergo Fenton-type reactions provoking the peroxidation of lysosomal membrane lipids, which results in its destabilization and permeabilization [32]. Subsequent release of lysosomal contents, including proteolytic enzymes to the cell cytoplasm, can potentially activate inflammatory and apoptotic pathways [31]. Moreover, lysosomal rupture will also severely impair the ability of the cell to dispose of misfolded proteins and aggregates.

Given the concurrence of oxidative stress, intermediate filament dysfunction and proteostasis alterations in AxD, here we have employed an astrocytoma cell model to assess whether the expression of AxD GFAP mutants is sufficient to elicit lysosomal alterations. Moreover, we have used this model to explore the vulnerability of these organelles to various insults, and in particular to oxidative stress.

## Material and methods

### Plasmids and transfections

The expression vector encoding the human GFAP ORF in fusion with GFP (GFP-GFAP) was from Genecopoeia. The bicistronic expression vector RFP//GFAP wt, coding for DsRed-Express2 and GFAP wt as separate products has been previously described [9]. The GFP-GFAP R239C mutation was introduced by site-directed mutagenesis as previously reported [9]. The point mutations R416W and the out-of-frame mutation D417M14X (DX; 1247-1249GGG>GG) were introduced into the GFP-GFAP plasmid by using the Quickchange XL Site-directed Mutagenesis kit (Agilent) or the NZYMutagenesis kit (NZYtech), using primers including the desired mutations (underlined), as follows: R426W forward, 5’-GGTGAAGACCGTGGAGATG<u>T</u>GGGATGGAGAGG-3’; DX forward, 5’-GGTGAAGACCGTGGAGATGCGG_ATGGAGAGGTCATTAAGG-3’), and their corresponding complementary reverse primers. Mutations were confirmed by DNA sequencing at Secugen S.L. (Madrid). The endolysosomal pH reporter plasmid mCherry-SEpHluorin-8 (pHluorin-8) has been previously described [33]. All plasmids used for transfection were purified using Endofree maxiprep kits (NZYtech). Transfections were carried out at 80 % cell confluence using Lipofectamine 2000 (Invitrogen), following the instructions of the manufacturer. Cells were incubated with transfection mixtures for 5 h in antibiotic-free medium. Experiments in transiently transfected cells were performed 48 h later. Selection of stable transfectants was achieved using cell culture media containing 500 μg/ml geneticin (G418, Gibco), for at least five passages after transfection, after which, expression of the construct of interest was confirmed by fluorescence microscopy. Enrichment of the cell population expressing GFP-GFAP was achieved by cell sorting on a cell Sorter FACSAria Fusion.

### Cell culture and treatments

U-87 MG astrocytoma cells were authenticated by microsatellite sequencing amplification (short tandem repeat (STR)-PCR profiling) at Secugen, S.L. (Madrid, Spain). Cells were grown in DMEM with 10 % (v/v) FBS (Gibco), 100 U/ml penicillin and 100 μg/ml streptomycin at 37°C in a humidified atmosphere with 5 % (v/v) CO_2_, and routinely checked for mycoplasma contamination at the Cell Culture Facility (CIB Margarita Salas, CSIC, Madrid). Treatment with H_2_O_2_ (Santa Cruz Biotechnology) was carried out at a final concentration of 1 mM in serum free medium for 30 min. LLOMe (CAS 6491-83-4, Santa Cruz Biotechnology) was used at a final concentration of 2 mM in growth medium with FBS and antibiotics.

### Immunofluorescence and confocal microscopy

Cells were grown on glass-bottom dishes (Mattek Corporation), glass coverslips or IbiTreat 5 mm imaging dishes with a polymer coverslip (Ibidi, 81156) and were fixed with 4 % (w/v) paraformaldehyde (PFA) in PBS for 25 min. Permeabilization with 0.1 % (v/v) Triton X-100 in PBS was done for 20 min, after which, blocking was performed with 1 % (w/v) BSA in PBS for 1 h at room temperature. Primary antibodies, anti-LAMP-1 (H4A3) (sc-20011, Santa Cruz), anti-galectin-1 (H-45) (sc-28248, Santa Cruz) and anti-galectin-3 (B2C10) (sc-32790, Santa Cruz) were used at 1:100 (v/v) dilution in blocking solution. After three washes with PBS, secondary antibodies conjugated to Alexa-488, Alexa-568 or Alexa-647 (Invitrogen), were used at 1:200 (v/v) dilution in the same buffer. In both cases, incubations were carried out for 1h at room temperature. Nuclei were counterstained with 2.5 μg/ml DAPI in PBS for 15 min. When indicated, lysosomes were stained with a 500 nM final concentration of LysoTracker™ Red DND-99 (LTR) (L7528, Thermo Fisher) for 20-30 min in serum free medium at 37 °C. For lysosomal staining of cells treated with H_2_O_2_, LTR and the oxidant were added at the same time. Cells were then fixed with 4 % (w/v) PFA before direct imaging or immunofluorescence, as indicated.

Cells were imaged on Leica SP5 or SP8 confocal microscopes with the 63x objective. Sections were acquired every 0.5 μm. Fluorescence intensity was adjusted below saturation, as determined with the LUT command. In assays requiring quantification of signal intensities, acquisition parameters were maintained between samples. In experiments performed to assess colocalization or galectin-1 and 3 internalization into lysosomes, images were acquired with the Lightning module of the Leica SP8 microscope and/or processed with the Lightning module after acquisition.

### Cathepsin B activity assay

Cathepsin B activity was measured using the Magic Red™ Cathepsin B Kit (ICT937, BioRad). The substrate was diluted 1:10 (v/v) in distilled H_2_O and then 1:25 (v/v) in serum free medium before addition. After 1 h, the medium was changed to DMEM with 10 % (v/v) FBS (Gibco), 100 U/ml penicillin and 100 μg/ml streptomycin. Live cells were imaged 20 h later.

### Image Analysis

Analysis and quantification of confocal images were done using FIJI 2.14.0 software. The entire cell area or the area of interest was labelled as a Region of interest (ROI), by manual selection or by use of the “Wand” tool in total projections. Fluorescence intensities (mean intensity of the ROI) were obtained with the “Measure” tool. Lysosomes were analyzed with the “Analyze particles” tool, considering LTR-stained particles with an intensity higher than 70 using the threshold tool, and with sizes larger than 0.1 µm^2^, in order to exclude the background noise. For estimation of the proportion of perinuclear lysosomes, a ROI was created by selecting the nuclear contour and enlarging the selection outwards to obtain a ring-shaped region 6 μm wide. For quantitation of the number of lysosomes in recovery experiments the “Threshold” tool was used, as above. When aggregates or collections of lysosomes were observed, the number of lysosomes within these collections was estimated by dividing their area by 0.5 µm^2^, since approximately 70% of the lysosomes in a healthy cell are below this size. The number of galectin-1 or galectin-3 puncta was quantified with the “Analyze particles” plugin, considering sizes above 0.02 µm^2^, in order to exclude the background noise. Colocalization analysis was performed with the BIOP JACoP plugin, using single sections from the different channels, acquired at mid cell height, and applying manual adjustment of the threshold in each ROI so the limits of the GFAP fibers and the lysosomes were correctly represented by the resulting mask created by BIOP JACoP for the analysis.

### Statistical analysis

Experimental data were collected from at least three independent experiments. Statistical analysis was carried out using Prism v9 8 (GraphPad), and results were shown as mean values ± standard error of the mean (SEM). Statistical differences between selected pairs of data sets were assessed using unpaired two-tailed Student’s *t*-tests and differences were considered significant when p < 0.05.

## Results

### Lysosomal distribution is altered as a result of GFAP mutation

We have previously reported that expression of certain GFAP AxD mutants is sufficient to alter organelle homeostasis, e.g. mitochondrial morphology and function, in astrocytic cells [9]. Here, we have used the U-87 MG astrocytoma cell line, which expresses very low levels of endogenous GFAP [9], to assess the consequences of the expression of GFAP wt or the AxD mutants R239C, R416W and D417M14X (DX) on lysosomal distribution and function. Expression of the corresponding constructs as GFP-GFAP fusion proteins in these cells elicits the formation of networks, constituted by nearly homogeneous populations of GFAP wt or the various mutants, which show distinct morphological patterns. Whereas the GFP-GFAP wt network appears formed by regular filaments, networks formed by the mutant proteins display diverse alterations ranging from a diffuse pattern to the presence of protein bundles and aggregates, characteristic of AxD, of different abundance (Fig. 1A). Coexistence of thick bundles and aggregates was particularly obvious in cells expressing GFP-GFAP R239C, as we previously reported [9].

**Figure 1.**
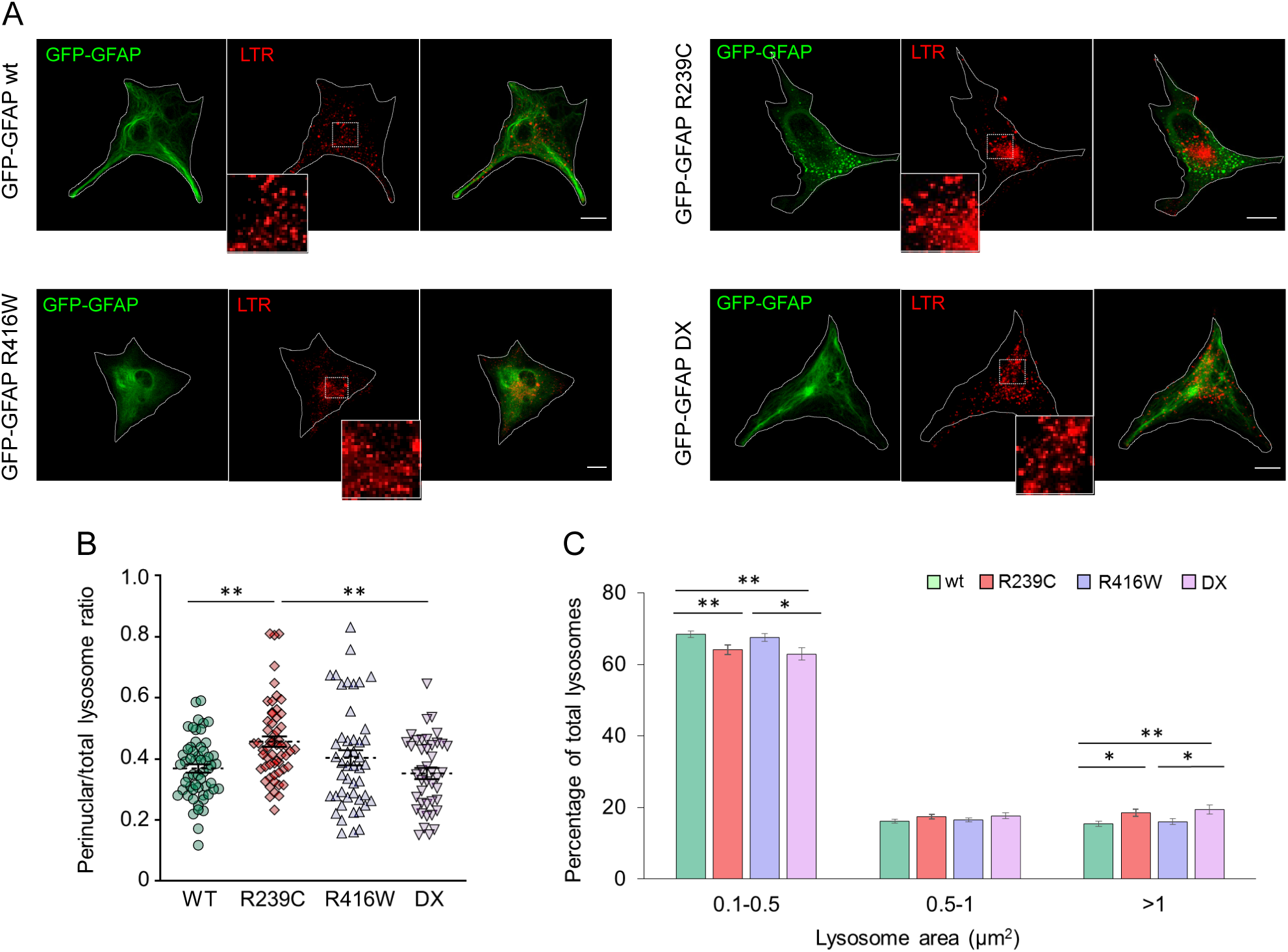
Lysosomal distribution in U-87 MG cells expressing GFP-GFAP wt or AxD mutants: (A) U-87 MG cells transiently transfected with GFP-GFAP (wt, R239C, R416W or DX mutants) were stained with Lysotracker Red (LTR). The distribution of GFP-GFAP and lysosomes was visualized by confocal microscopy. Cell contours are indicated by dotted lines. Insets show enlarged areas of interest. (B) Proportion of lysosomes located in the perinuclear area vs total lysosomes, defined as detailed in Methods. (C) Size distribution of lysosomes. The area of lysosomes was measured with FIJI and the proportion of particles within the indicated categories was quantitated. Particles with size over 1 μm^2^ include large lysosomes and groups of several lysosomes, which could not be fully individualized. Results shown are average values ± SEM of at least 47 (B) or 63 determinations (C), from at least 4 separate experiments *p<0.05; **p<0.01. Scale bars, 20 μm.

The distribution of lysosomes in cells expressing GFAP wt or mutants was monitored by staining with LysoTracker™ Red (LTR), which labels acidic compartments within the cell. Cells expressing GFP-GFAP wt displayed a typical lysosomal localization, with a nearly homogeneous distribution throughout the cytoplasm, and a partial perinuclear enrichment (Fig. 1A). Conversely, expression of GFP-GFAP mutants elicited diverse alterations in lysosomal position and appearance. Thus, cells expressing GFP-GFAP R239C frequently showed a juxtanuclear accumulation of lysosomes (Fig. 1A), which displayed a larger size and/or appeared in dense groups (Fig. 1B and 1C). GFP-GFAP R416W did not alter the average lysosomal size (Fig. 1C), although approximately 25% of cells displayed a preferential perinuclear distribution of lysosomes. In turn, GFP-GFAP DX led to an increase in the size of lysosomes and/or the number of aggregates (Fig 1C), without apparently affecting their distribution (Fig. 1B).

As shown above, GFP-GFAP R239C elicited marked alterations in both the localization and size of lysosomes in U-87 MG astrocytoma cells. Therefore, we explored the impact of the GFAP R239C mutation on lysosomal distribution in more detail. First, we evaluated the position of lysosomes with respect to GFAP filaments (Fig. 2). In cells expressing GFP-GFAP wt most lysosomes appeared located in the spaces between GFAP filaments or associating laterally with them (Fig. 2A). In cells expressing GFP-GFAP R239C, lysosomes also associated laterally with GFAP filaments showing a degree of overlap similar to that found in cells expressing GFP-GFAP wt, but were mostly excluded from GFAP R239C bundles and aggregates (Fig. 2A). This association was quantitated by Manders coefficient, which rendered 33% and 37% overlap of LTR over GFP-GFAP wt and GFP-GFAP R239C, respectively. Next, we confirmed these observations by monitoring the distribution of the protein LAMP-1, which is present in late endosomes and lysosomes. LAMP-1 positive vesicles appeared homogeneously distributed in cells bearing GFP-GFAP wt, showing a lateral association with GFAP wt filaments (Fig. 2B). In contrast, cells expressing GFP-GFAP R239C display a marked perinuclear accumulation of LAMP-1 structures, consistent with the altered lysosomal distribution described above. Moreover, a lateral coincidence of LAMP-1 and GFP-GFAP R239C signals was apparent, and as with LTR, the presence of LAMP-1 very close to GFAP R239C bundles could be appreciated. Nevertheless, LAMP-1 positive structures were very scarce within the bundles. Manders’ coefficient analysis showed a 59% overlap between LAMP-1 and GFP-GFAP wt signals and a 50% overlap for GFP-GFAP R239C.

**Figure 2.**
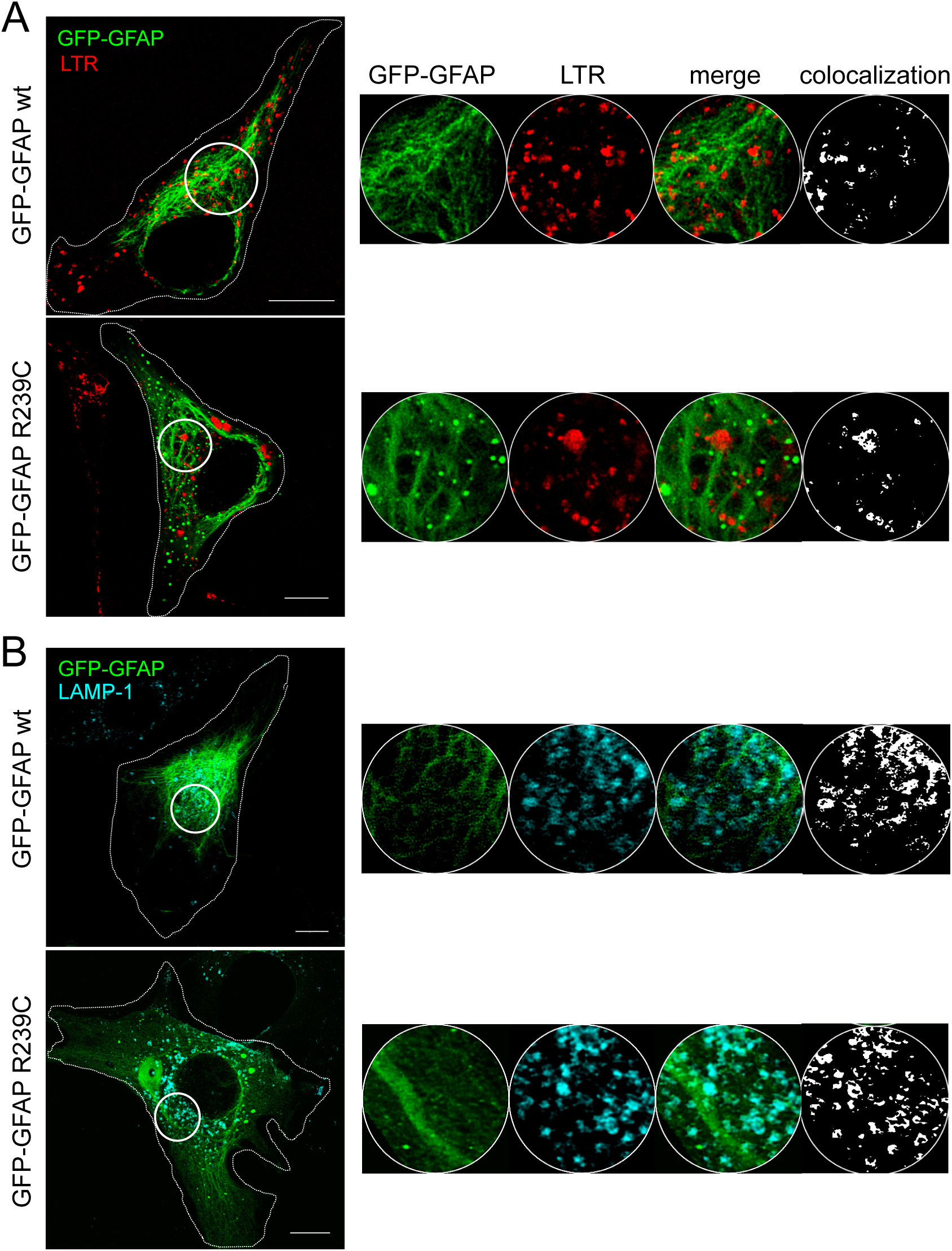
Relative positions of lysosomes and GFAP filaments: U-87 MG cells expressing GFP-GFAP wt or R239C were stained with LTR (A) or immunostained with anti-LAMP-1 (B). GFP-GFAP is depicted in green and lysosomes in red (LTR) or cyan (LAMP-1). Overall projections are shown. The areas delimited by the white circles are enlarged in the right panels showing single sections of the individual and merged channels, as well as colocalization masks for GFAP and lysosomes (far right). Scale bars, 10 µm.

### Lysosomal function is altered in U-87 MG astrocytoma cells expressing the AxD mutant GFP-GFAP R239C

In view of the anomalies in the distribution of lysosomes encountered in cells bearing the GFP-GFAP R239C mutant, we next explored putative alterations in their function. First, we assessed the activity of cathepsin B, one of the main lysosomal proteases, using the Magic Red™ Cathepsin B Kit (Fig. 3A). In this assay, cleavage of a cathepsin B substrate releases a cresyl violet fluorophore, the intensity of which provides an index of the enzymés activity. Interestingly, the Magic Red signal yielded the typical lysosomal pattern described above for cells expressing GFP-GFAP wt or the R239C mutant. Nevertheless, the intensity of the signal detected in cells expressing GFP-GFAP R239C was significantly lower than that of cells expressing GFP-GFAP wt (Fig. 3B), indicating a marked decrease in lysosomal cathepsin B activity in cells bearing the AxD GFAP mutant. Moreover, the proportion of the cell surface covered by detectable Magic Red fluorescence was also smaller in cells expressing GFP-GFAP R239C than in those carrying GFP-GFAP wt (Fig. 3C). These results indicate that in GFAP R239C cells there is a decrease in the global cathepsin B activity, as well as in the number of lysosomes with detectable cathepsin B activity, which reveals an impairment in their lysosomal function.

**Figure 3.**
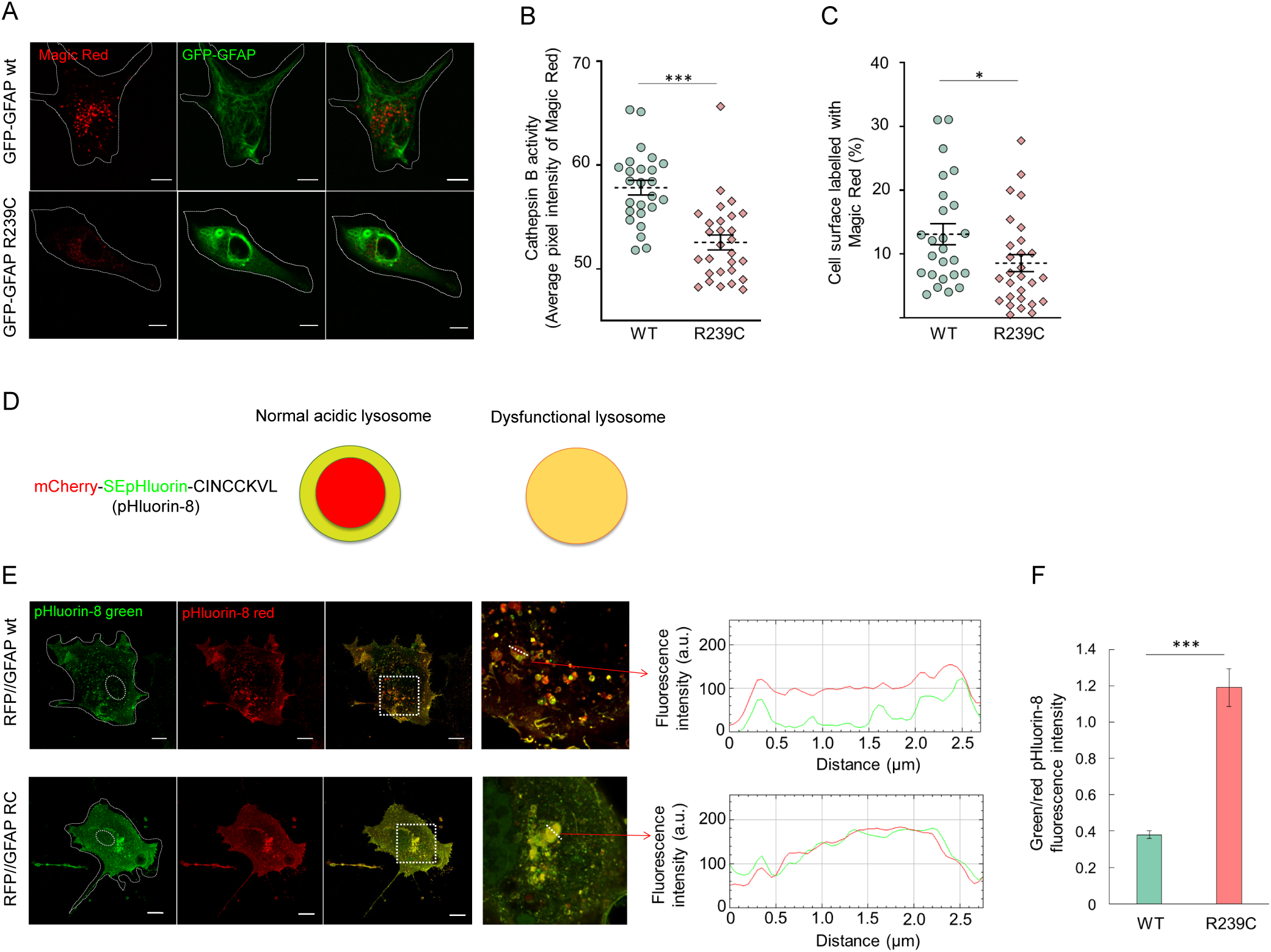
Evaluation of lysosomal function in cells expressing GFP-GFAP wt or R239C. U-87 MG cells transfected with the indicated GFP-GFAP plasmids were incubated with Magic Red Cathepsin B substrate, which emits a fluorescent signal when cleaved by Cathepsin B, and live cells were visualized by confocal microscopy (A). Overall projections of individual and merged channels are shown. Scale bars, 10 μm. Cathepsin B activity, estimated as the average pixel intensity (B) and proportion of the cell covered by the fluorescence of the Magic Red cleaved substrate (C) were quantitated by analysis of images from three independent experiments totaling a minimum of 20 cells. Results shown are average values ± SEM. *p<0.05; ***p<0.001. Scale bars, 10 μm. (D) Schematic representation of the mCherry-SE-pHluorin-CINCCKVL (“pHluorin-8”) construct, and the expected signal in healthy, acidic lysosomes (selective loss of the green signal inside the lysosome), and in dysfunctional lysosomes (preservation of both, red and green signals inside the lysosome). (E) U-87 MG cells were transiently cotransfected with RFP//GFAP wt or R239C and pHluorin-8 to assess intralysosomal pH by monitoring the loss of pHluorin-8 green fluorescence inside lysosomes. Images shown are single sections taken at mid-cell height. Regions of interest delimited by the squares are enlarged at the right. Fluorescence intensity profiles of the pHluorin-8 green and red signals along the dashed lines drawn across single lysosomes are depicted as indicated by the red arrows. Scale bars, 10 μm. (F) The ratio of green to red fluorescent signals at the mid-point of the lysosome was calculated from at least 5 lysosomes per cell and 10 cells per experiment, and is shown as average values ± SEM. ***p<0.001.

The maintenance of an acidic intraluminal pH is necessary for the optimal activity of lysosomal proteases. Therefore, next we monitored intralysosomal pH by means of the pH sensitive reporter mCherry-SEpHluorin-8 (pHluorin-8) [33]. This construct encodes a pH insensitive fluorescent protein (mCherry), fused to a pH sensitive fluorescent protein (SEpHluorin), bearing an endolysosomal localization signal at its C-terminal end (Fig. 3D). This peptide signal (CINCCKVL, abbreviated as “8”) corresponds to the C-terminus of the small GTPase RhoB [34], and it has been previously confirmed to direct the endolysosomal localization of chimeric proteins [35, 36]. In neutral environments, both the red and green fluorescent signals are detected, whereas in acidic environments there is a loss of the green fluorescence [33]. In these assays, we transfected astrocytoma cells with vectors encoding untagged GFAP variants in order to avoid overlap of fluorescent tags with the signals from the pHluorin-8 reporter. Interestingly, in cells expressing GFAP wt, pHluorin-8 appeared as small dots and vesicles distributed throughout the cytoplasm (Fig. 3E, upper images). Magnification of selected areas evidenced the characteristic pattern of pHluorin-8 fluorescence when expressed in combination with GFAP wt. In particular, the lysosomal contour exhibited both red and green fluorescence, whereas the lysosomal lumen exhibited mainly red fluorescence, indicative of an acidic intraluminal pH. In sharp contrast, astrocytoma cells expressing GFAP R239C exhibited a diffuse pHluorin-8 cytoplasmic background together with an accumulation of the pH reporter in large endolysosomes near the nucleus, that showed a complete overlap of red and green signals of similar intensities (Fig. 3E, lower images). These differences are clearly illustrated in the corresponding fluorescence intensity profiles (Fig. 3E, right), which showed the typical decline of green fluorescence in the lumen of endolysosomes from cells transfected with GFAP wt, whereas no decrease of the green signal occurred in vesicles from GFAP R239C expressing cells, indicating a loss of their acidic intraluminal pH. Indeed, quantitation of this effect showed that the ratio of intraluminal green/red fluorescence intensities of pHluorin-8 was three times higher in cells expressing GFAP R239C (Fig. 3F). Taken together, these results reveal a severe compromise of lysosomal function in cells expressing the AxD GFAP mutant, characterized by impaired lysosomal acidification and decreased cathepsin B activity.

### U-87 MG astrocytoma cells expressing the GFP-GFAP R239C AxD mutant are more vulnerable to lysosomal damage induced by oxidative stress

We have previously reported that astrocytoma cells expressing GFP-GFAP R239C are more susceptible to oxidation and lipoxidation than those expressing the wt protein [9]. Indeed, exposure to oxidants, such as H_2_O_2_, or to electrophilic lipids, such as hydroxynonenal (HNE) or 15-deoxy-protaglandin J_2_ (15d-PGJ_2_), elicits an increase in GFAP R239C aggregates [9]. Therefore, we next explored whether expression of the AxD GFP-GFAP R239C mutant affected the susceptibility of lysosomes to oxidative stress (Fig. 4). First, we established the conditions to elicit a detectable lysosomal damage, by treating cells with a bolus administration of H_2_O_2._ Incubation with 1 mM H_2_O_2_ for 30 min significantly decreased LTR staining, indicating a loss of detectable lysosomes in cells expressing either the wt or the R239C protein (Fig. 4A). Notably, approximately 10% of the cells displayed an increase in a diffuse red cellular background, indicative of lysosomal leakage (Suppl. Fig. 1, lower panels). This damage is clearly illustrated in the fluorescence intensity profiles (Suppl. Fig. 1, right panels). To quantitate the extent of the damage in cells expressing GFP-GFAP wt or R239C, several parameters were employed. The number of lysosomes, defined as LTR-positive particles, was quantitated as an index of their abundance in each cell (Fig. 4B). The average loss of LTR-positive particles was more severe in cells expressing GFP-GFAP R239C (60%) than in those carrying the wt protein (33%), a significant difference that points to a higher susceptibility of lysosomes to oxidative stress in cells expressing the AxD mutant (Fig. 4B). In addition, lysosomal compartments were monitored by immunofluorescence with anti-LAMP-1 antibody (Fig. 4C). Treatment with H_2_O_2_ did not significantly affect the average intensity of the LAMP-1 signal in cells expressing GFP-GFAP wt (Fig. 4D). However, in cells expressing GFP-GFAP R239C H_2_O_2_ elicited a significant decrease in LAMP-1 intensity, suggestive of lysosomal loss (Fig. 4D). Taken together, these results indicate a greater lysosomal fragility in cells expressing the AxD GFAP mutant upon exposure to H_2_O_2_.

**Figure 4.**
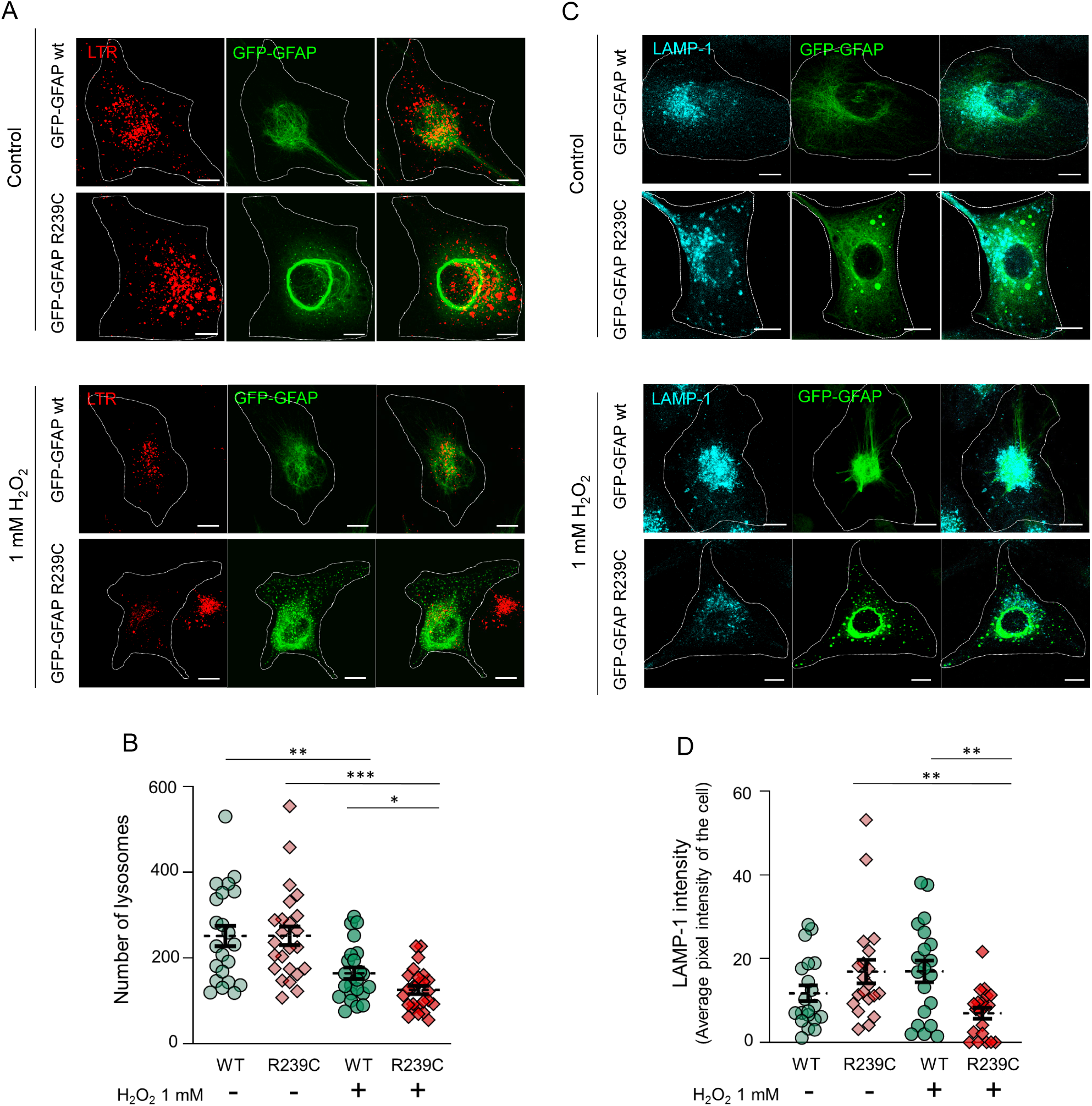
Effect of H_2_O_2_ on LTR and LAMP-1 signal intensities in U-87 MG cells expressing GFP-GFAP wt or R239C. Cells were treated with vehicle or 1 mM H_2_O_2_ for 30 min. For LTR staining (A), the dye was included during this incubation. For LAMP-1 detection (C), cells were fixed after treatment and subjected to immunofluorescence with anti-LAMP-1. Overall projections of single and merged channels are shown. The number of detectable lysosomes (B) and the intensity of the LAMP-1 signal (D) were quantitated in images from three independent experiments totaling a minimum of 20 cells. In the case of lysosomal accumulations, the number of lysosomes within these gatherings was estimated as the ratio between the total area of the conglomerate and an estimated lysosomal size of 0.5 μm^2^. Results are shown as average values ± SEM. *p<0.05; **p<0.01; ***p<0.001. Scale bars, 10 μm.

Extensive lysosomal damage is often accompanied by permeabilization or rupture of the lysosomal membrane. Under these conditions, galectins are recruited to the lysosomal interior, where they bind the sugar moieties of the glycoproteins that line the inner side of the lysosomal membrane (schematized in Fig. 5A). This may be followed by the onset of lysosomal repair mechanisms and/or recruitment of ubiquitin ligases that target the damaged lysosomes for degradation [37, 38]. In order to assess whether these mechanisms were in place in the astrocytoma cellular model, we performed immunodetection of galectin-1 and galectin-3 in cells expressing GFP-GFAP wt under control conditions and after treatment with H_2_O_2_, as above (Fig. 5B and C). This staining allows the identification of damaged lysosomes as sites of galectin’s accumulation (galectin “puncta”). Galectin-1 or -3 puncta were barely detectable in control cells, but became obvious after H_2_O_2_ treatment. Interestingly, in some cases, colocalization of the LTR signal with particulate galectin staining could be observed, which likely corresponds to damaged lysosomes that have not lost all their contents and acidity (Fig. 5B and C, right images). It should be noted that in these experiments LTR staining was partially lost during permeabilization and immunofluorescence, which explains the low levels observed. Next, we employed this assay to characterize lysosomal damage in cells expressing GFP-GFAP wt or R239C. We observed that under control conditions, galectin-1 puncta were more abundant (average of 20 puncta/cell) in cells expressing the AxD GFAP mutant (Fig. 5D and E), which could indicate a potential “constitutive” lysosomal damage or increased lysosomal fragility. Treatment with H_2_O_2_ increased the number of galectin-1 puncta in both GFAP wt and R239C expressing cells (Fig. 5E). Nevertheless, galectin-1 puncta were significantly more abundant in cells expressing GFP-GFAP R239C, with approximately 50% of the cells displaying more than 50 galectin-1 puncta/cell. Detection of galectin-3 yielded a similar tendency, although in this case, the number of galectin-3 puncta was generally lower, and only increased significantly upon treatment of cells expressing GFP-GFAP R239C with H_2_O_2_ (Fig. 5F and G). Images of single and merged channels, including LTR staining, are provided in Suppl. Fig. 2. Altogether, these results indicate that lysosomes of cells expressing the AxD GFP-GFAP R239C mutant display increased fragility and are more susceptible to damage by oxidative stress than those present in cells expressing GFP-GFAP wt. Moreover, the intense galectin staining indicates that the damage of these organelles may be severe.

**Figure 5.**
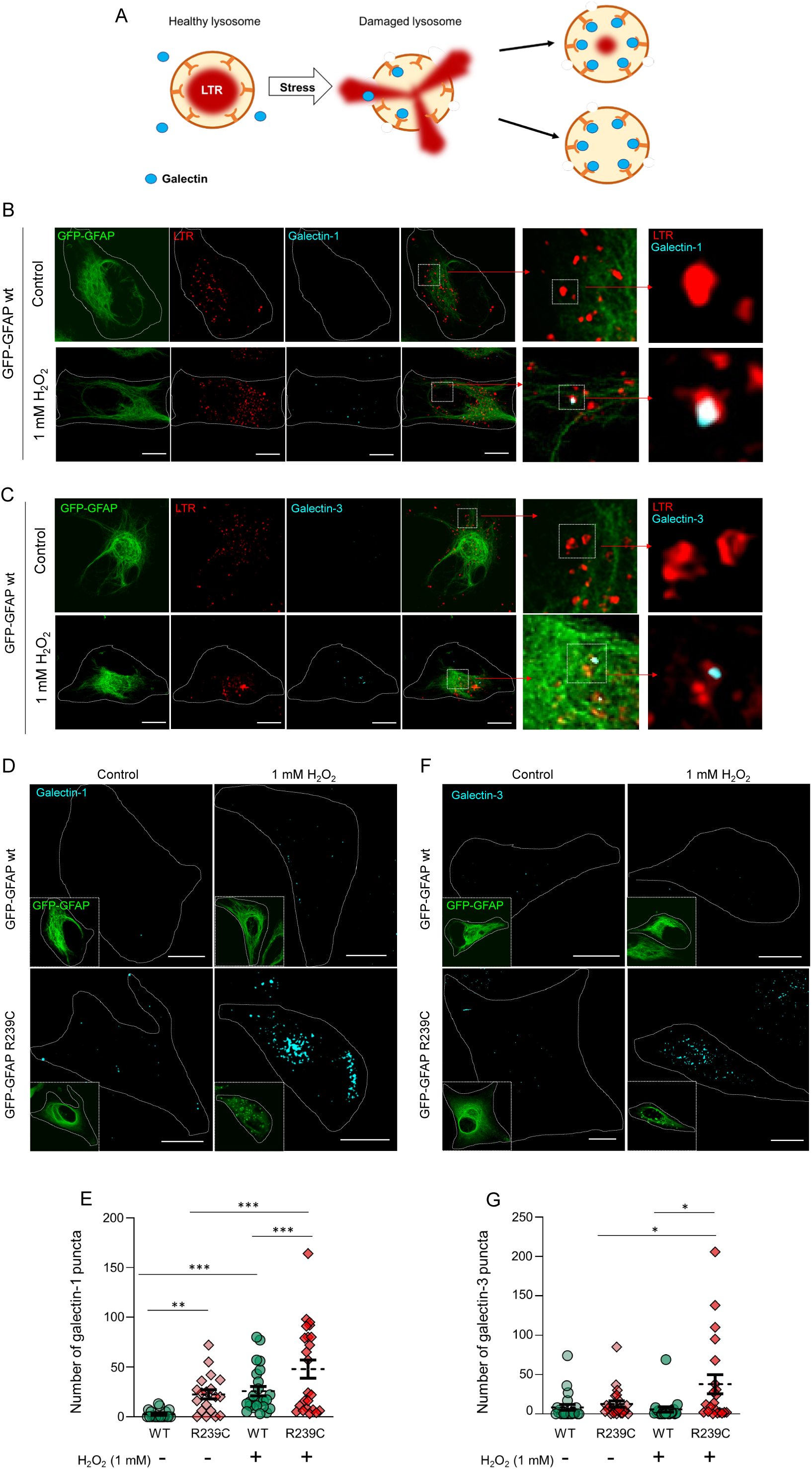
Detection of galectins in damaged lysosomes. (A) Scheme of lysosomal damage releasing lysosomal contents and allowing galectins to bind to sugars lining the internal lysosomal surface. (B) and (C) Detection of galectin puncta in association with LTR positive particles in damaged lysosomes. U-87 MG cells expressing GFP-GFAP wt were treated with H_2_O_2_, as above and stained with LTR and antibodies against galectin-1 (B) or galectin-3 (C). Single channels and merged images are shown. Areas of interest, delimited by dotted squares are successively enlarged at the right. Far right images illustrate colocalization of galectins and LTR signals in the cells treated with H_2_O_2_, indicative of galectin recruitment into damaged lysosomes, which is not found in the control cells. (D-G) Galectin-1 and Galectin-3 immunostaining of cells subjected to oxidative stress. U-87 MG cells expressing GFP-GFAP wt or R239C were treated with vehicle or 1 mM H_2_O_2_ for 30 min and stained with LTR (shown in Suppl. Fig. 2) and antibodies against galectin-1(D) or galectin-3 (F) (cyan). GFAP-GFAP fluorescence is shown in insets. The number of galectin puncta per cell, indicative of lysosomal recruitment, was quantitated by image analysis as described in Methods and depicted in (E, galectin-1) and (G, galectin-3). Results shown are average values ± SEM of a total of at least 20 cells from 3 independent experiments. *p<0.05; **p<0.01; ***p<0.001. Scale bars, 10 μm.

To confirm the defective condition of lysosomes in the AxD cellular model, we explored their response to H-Leu-Leu-OMe hydrochloride (LLOMe), a compound that is converted into membranolytic metabolites in lysosomes causing lysosomal rupture [39]. Notably, this compound induced cell rounding and loss of adherence in U-87 MG cells cultured on glass coverslips. Therefore, to minimize cell detachment, for these assays, cells were seeded on Ibidi-treated cell culture dishes. In this experimental setting, cells expressing the AxD GFP-GFAP R239C mutant also tended to display more galectin-1 and galectin-3 puncta under control conditions (Fig. 6). Remarkably, exposure to LLOMe elicited clear lysosomal damage, demonstrated by the increase in the number galectin-1 and galectin-3 positive particles (or puncta), which was again more evident in cells expressing GFP-GFAP R239C (quantitated in Fig. 6B and D). These results confirm that in this cellular model, expression of the AxD GFP-GFAP R239C mutant leads to increased lysosomal fragility.

**Figure 6.**
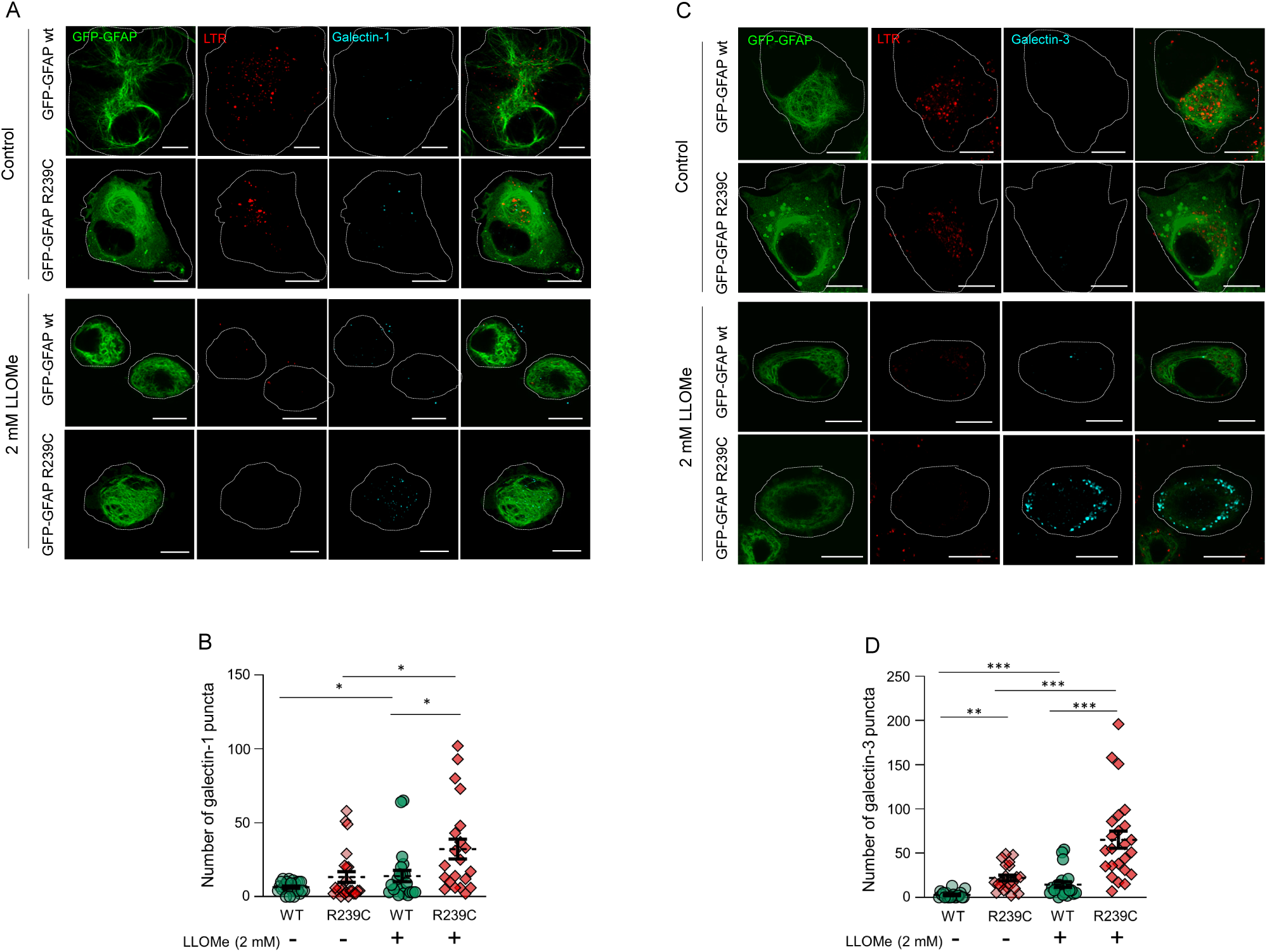
Galectin-1 and Galectin-3 immunostaining in cells U-87 MG treated with LLOMe. U-87 MG cells expressing GFP-GFAP wt or R239C (A and C) were treated with vehicle or 2 mM LLOMe for 30 min and stained with LTR (red) and (A) anti-galectin-1 or (B) anti-galectin-3 (cyan). Individual channels and merged images are shown. The number of galectin puncta per cell was assessed by image analysis in a total of at least 20 cells from 3 independent experiments (B, D). Results are shown as average values ± SEM. *p<0.05; **p<0.01; ***p<0.001. Scale bars, 10 μm.

### Lysosomal repair is defective in U-87 MG GFP-GFAP R239C cells

In case of lysosomal damage, several cellular repair mechanisms can be activated to restore lysosomal number and function. Whereas limited damage can be overcome by repair of the lysosomal membrane, more severe damage requires clearance of affected lysosomes and generation of new ones [38, 40]. Importantly, galectins participate in the coordination of these repair pathways [41]. Hence, given the increased recruitment of galectins to damaged lysosomes in cells expressing GFP-GFAP R239C, we next attempted to detect signs of lysosomal repair (Fig. 7). For this purpose, cells were treated with H_2_O_2_ for 30 min, as above, after which, the oxidant was removed and cells were allowed to recover for 1 h or 4 h. Consistent with the results shown in Fig. 4, H_2_O_2_ treatment elicited a decline in the number of LTR positive particles, indicative of lysosomal damage, which was more intense in cells expressing the AxD GFP-GFAP R239C mutant. Interestingly, upon removal of the stressor, cells bearing GFP-GFAP wt showed a sharp increase in the number of lysosomes, which remained stable for several hours, indicating lysosomal recovery. In contrast, removal of the oxidant from cells expressing GFP-GFAP R239C resulted in a marginal increase in LTR positive vesicles, which did not attain the values of untreated cells even after 4 h (Fig. 7B). Single channels and overlay images of these results are provided in Suppl. Fig. 3. Together, these observations point to profound anomalies in the recovery of lysosomes from oxidative stress in cells expressing GFP-GFAP R239C, and suggest that AxD GFAP mutations may compromise lysosomal repair.

**Figure 7.**
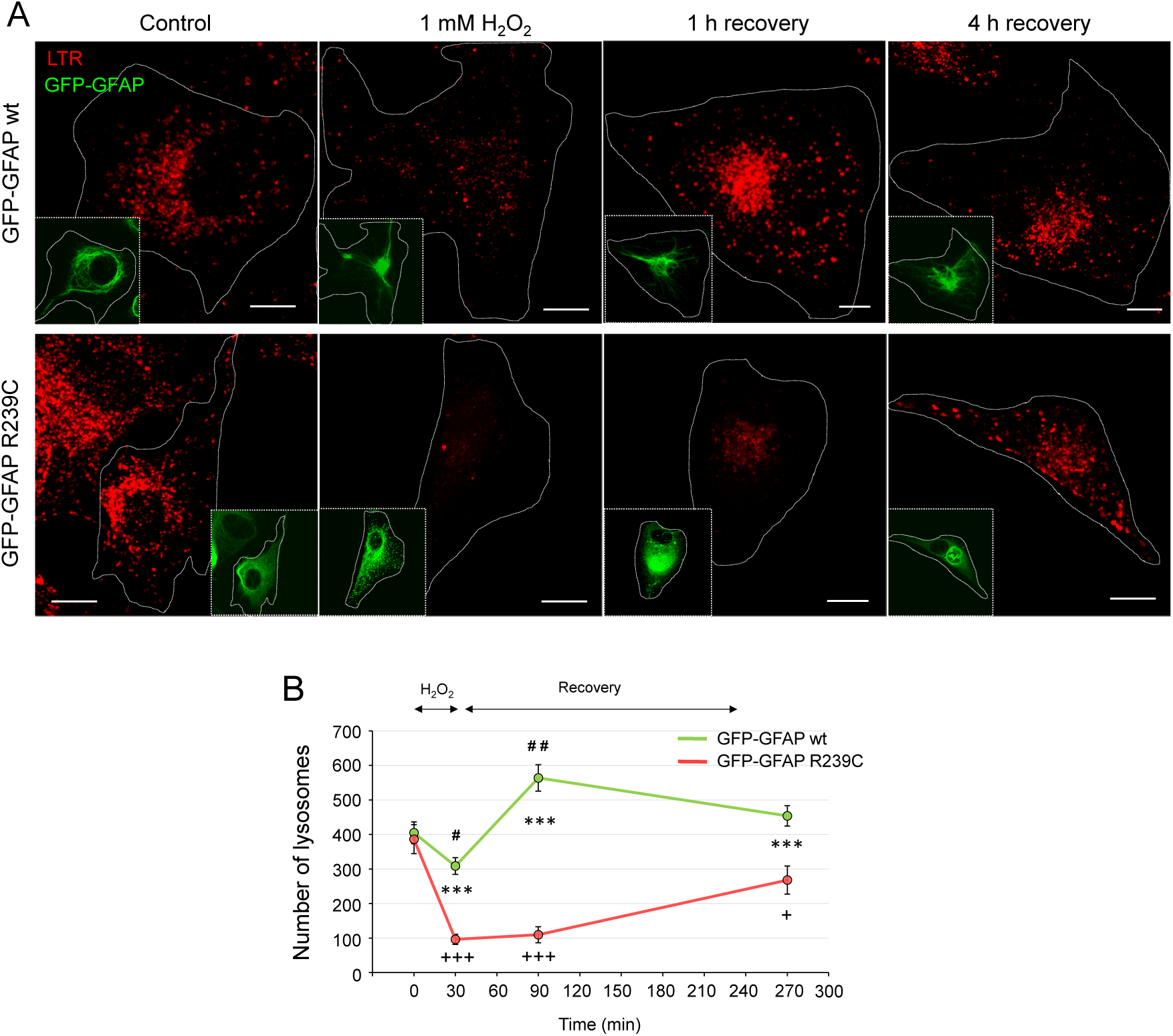
Assessment of lysosomal recovery after H_2_O_2_-induced damage. U-87 MG cells expressing GFP-GFAP wt or R239C were treated with 1 mM H_2_O_2_ for 30 min, after which the oxidant was removed and cells were allowed to recover for 1 h or 4 h in complete medium (A). LTR staining was carried out as described in the Methods section. At the indicated time points, cells were fixed and subsequently imaged by confocal microscopy. Overall projections of single channels are shown, depicting LTR fluorescence (main images), and GFP-GFAP (insets). (B) The number of lysosomes per cell, identified as LTR-positive vesicles, was obtained by image analysis. Results are average values ± SEM from 3 independent experiments totaling at least 20 cells. ***p<0.01 wt vs R239C for each time point; ^#^p<0.05, ^##^p<0.01 vs 0 time for wt; ^+^p<0.05; ^+++^p<0.001 vs 0 time for R239C. Scale bars, 10 μm.

## Discussion

In the rare leukodystrophy known as AxD, GFAP mutations are considered to provoke abnormal filament network formation and increased GFAP levels leading to a pathologic “gain of function” causing dysfunction of numerous cellular organelles and compartments, altered mechanosensing and impaired proteostasis, which compromise astrocytic functions in brain support, or even elicit toxic consequences, leading to neurodegeneration [42–44]. Although there have been significant advances in the understanding of AxD, there are multiple aspects that still need to be elucidated, including the precise connections between GFAP mutations, altered assembly and pathology. Unveiling the pathogenic sequence of events requires a characterization of the full extent of the changes elicited by GFAP mutants at the cellular level. Our previous work indicates that expression of GFAP in U-87 MG astrocytoma cells provides a robust model to explore the cellular pathology induced by GFAP mutations [9]. Using this model, we have evidenced the different impact of several GFAP mutations on filament assembly, mitochondrial morphology and oxidative stress. Moreover, in the case of the severe R239C mutation we have proposed a pathogenic cycle involving the generation of oxidative stress and increased oxidation of the mutant protein, driving and/or feeding from altered mitochondrial function and aggregate formation [9]. Lysosomes are key players and targets of oxidative stress [31]. Therefore, here we have explored the influence of GFAP mutants, and specially R239C in lysosomal morphology and function in the astrocytoma cell model.

Our results show that just the expression of several GFAP AxD mutants in this model is sufficient to alter lysosomal distribution and size. The harshest alterations, in the form of juxtanuclear accumulations and increased lysosomal size, were observed in cells expressing the R239C mutant, consistent with the severity of this mutation. Several mechanisms could be envisaged for this effect. Lysosomal position and size are regulated in part by endosomal GTPases. In particular, Rab7 interacts with dynein and kinesins to ensure that lysosomes are correctly transported along microtubules and reach their appropriate locations [45, 46]. Interestingly, interactions of GFAP and/or its partner vimentin with proteins along the endolysosomal pathway, including endosomal GTPases Rab5a, Rab7, the adaptor complex AP3 and the lysosomal protein Lamp2A have been described in various experimental models [30, 47–49]. In particular, absence of vimentin or expression of certain mutants has been reported to alter lysosomal localization, with impaired cytoplasmic distribution and juxtanuclear accumulation [29], whereas mislocalization of lysosomes has also been observed in astrocytes differentiated from AxD patient-derived iPSCs [8]. Interestingly, alterations in lysosomal localization and function have been associated with several kinds of stress such as starvation or pathogen infections [50, 51], as well as in other neurological diseases, such as Huntingtońs disease [52], which also presents dysregulation of proteostasis and abnormal aggregates.

Besides location, the size of lysosomes is tightly regulated, as it can affect lysosomal function [53]. Enlargement of the lysosomes has been correlated with functional alterations including reduced recruitment of hydrolases and acidification defects [53]. Interestingly, here we have observed an increase in the proportion of large lysosomes in cells expressing the GFAP R239C mutant, together with impaired cathepsin B activity and defective lysosomal acidification. All these alterations are clear indicators of lysosomal malfunction. Along with L and D, cathepsin B is one of the most abundant cathepsins in lysosomes. Both the proteolytic cleavage-dependent activation of pre-cathepsins inside lysosomes as well as the activity of the mature enzymes are greatly dependent on lysosomal pH, with optimal values around 4.5 [54, 55]. Therefore, the acidification defect is expected to have important consequences in the performance of lysosomes. Importantly, intermediate filaments, and in particular vimentin, were early reported to affect lysosomal pH since vimentin deficient fibroblasts showed decreased staining of organelles with the pH sensitive dye Lysosensor Green, compared to their wild type counterparts [30]. Interestingly, GFAP has been recently reported to interact with ATP6V1B2 [47], the regulatory subunit of vacuolar ATPase (V-ATPase), which is involved in the acidification of intracellular compartments [56]. Importantly, mutations in V-ATPase subunits or accessory proteins cause various familiar neurodegenerative diseases [57, 58].

As mentioned above, the expression of the AxD GFAP R239C mutant elicits oxidative stress, which poses a great threat to cellular organelles. Under oxidative stress, the lysosomal membrane becomes exposed to radicals arising from Fenton-type reactions that provoke its permeabilization [31, 32]. Indeed, in this work, in addition to a basal functional defect, we have observed a higher vulnerability of lysosomes of cells expressing GFAP R239C to both exogenously provoked oxidative stress and chemically induced rupture. Lysosomal damage is evidenced by the decrease in both detectable LTR-positive vesicles and LAMP-1 signal intensity. The decrease in LTR signal could be related to impaired acidification and/or to the leakage of lysosomal contents, suggested by the increased cytoplasmic LTR staining. In turn, a lower LAMP-1 signal, could reflect an actual loss of membrane components, which is more severe in GFAP R239C expressing cells. Importantly, release of lysosomal material to the cytoplasm, including enzymes and metals, would feed to the stress already present in the cell, and eventually lead to apoptosis or necrosis, hypothetically adopting the form of ferroptosis [31], and would compromise cell survival after stress [59]. Moreover, analogously to the pathogenic feedback loop arising from disruption of mitochondria, release of lysosomal contents could worsen GFAP assembly and contribute to the formation of aggregates, which in turn, would be deleterious for organelle homeostasis, thus closing a vicious circle, as schematized in Fig. 8.

**Figure 8.**
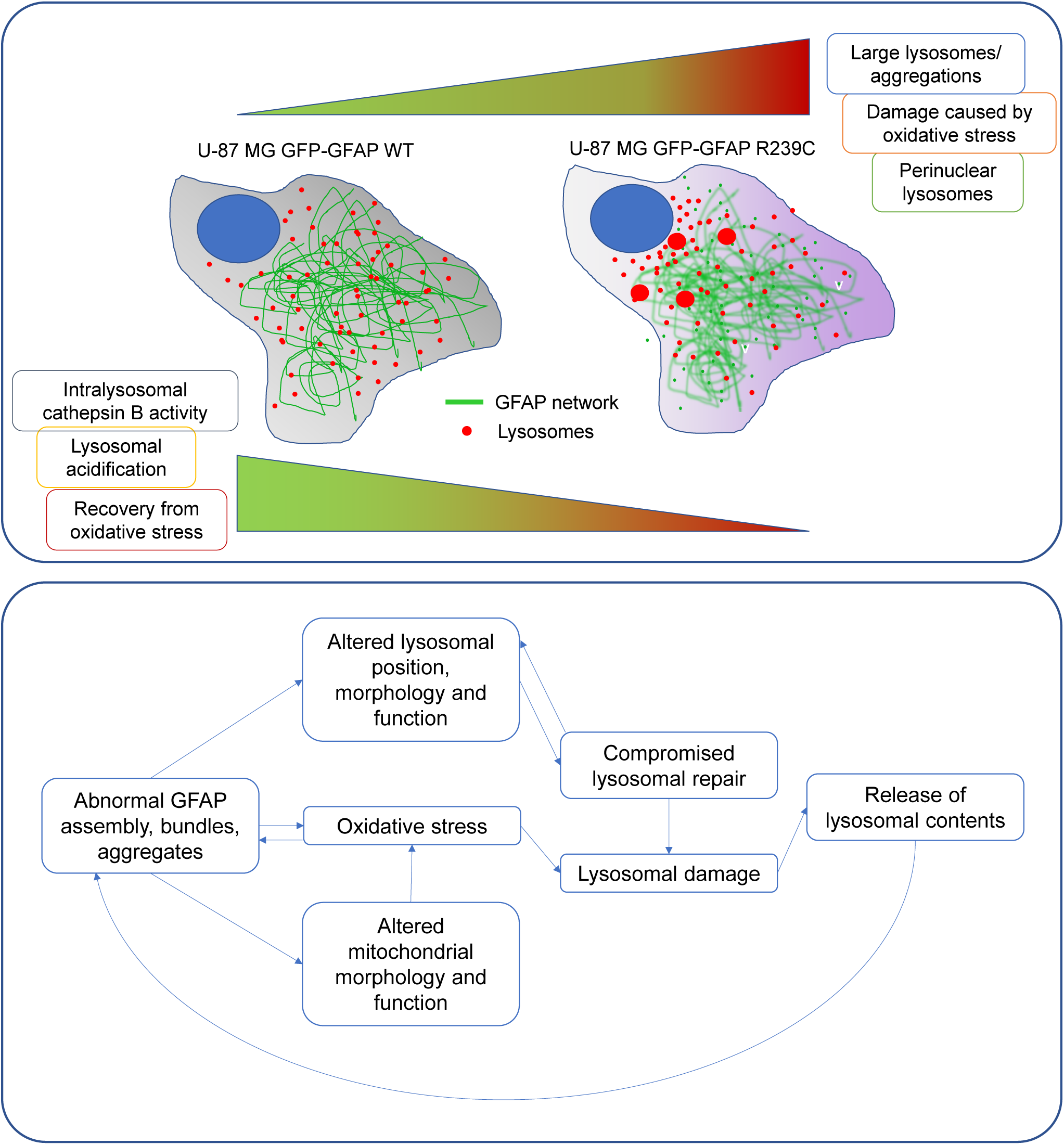
Summary of the findings of our study. Expression of the AxD mutant GFAP R239C alters lysosomal distribution, with perinuclear concentration and appearance of lysosomal clumps, as well as function, involving a defect in intracellular acidification, lower cathepsin B activity, increased fragility and compromised recovery after damage (upper panel). These effects, together with alterations of other organelles, could constitute a vicious circle amplifying the damage (lower panel).

Severe lysosomal damage exposes glycans of the inner leaflet of the lysosomal membrane that recruit galectins [41]. Several galectins, including galectin-1 and -3, can translocate inside damaged lysosomes, appearing as puncta, making it possible to label these organelles even before other signs of severe damage are evident [37]. Indeed, we have noticed that, under control conditions, cells bearing GFP-GFAP R239C tend to display more galectin puncta than those expressing GFP-GFAP wt, suggesting an increased fragility. Moreover, exposure of the cells to stressors such as H_2_O_2_ or LLOMe led to a clear increase in the number of galectin-1 and galectin-3 puncta, indicative of lysosomal damage that is more evident in cells expressing GFP-GFAP R239C. Some of the damaged lysosomes were co-stained with LTR and anti-galectin, illustrating the recruitment of galectins to lysosomes that still retain some of their contents. Of interest, galectins are also targets of oxidative and electrophilic modifications [60–63], which could affect lysosomal repair mechanisms under severe oxidative stress. Altogether, these results clearly show that lysosomes in cells expressing GFP-GFAP R239C are defective in integrity and function, and more vulnerable to ensuing oxidative or chemical stress, than those present in cells with GFP-GFAP wt.

Lysosomal damage can be exacerbated by the failure of repair mechanisms. Interestingly, we observed that cells expressing GFAP wt could attain an efficient lysosomal recovery upon removal of the oxidant, whereas the restoration of the number of acidic particles was blunted in cells expressing the AxD GFAP R239C mutant within the time frame of the assay. Galectins are more than markers of lysosomal damage and accumulating evidence indicates that they orchestrate lysosomal repair [41]. Galectin-3 internalization into the lysosome has been reported to trigger several pathways promoting lysosomal repair and induction of lysosomal biogenesis. In lysosomes with mild damage, galectin-3 facilitates repair by recruiting the Endosomal Sorting Complex Required for Transport (ESCRT) machinery, which can reseal small fissures in the membrane [41, 64]. If the damage is more severe, galectin-3 can also activate autophagy for removal of damaged lysosomes (lysophagy) and/or lysosomal biogenesis for replacement [41]. The dynamic balance of galectins during lysosomal damage also influences TFEB [41], a master regulator of lysosomal biogenesis and repair that binds to the promoters of autophagy genes involved in autophagosome biogenesis and autophagosome–lysosome fusion [65]. TFEB also promotes the generation of lysosomes and the expression of lysosomal enzymes; therefore, its activation should result in the substitution of the damaged lysosomes by new ones [66]. Our results suggest that lysosomal repair mechanisms may be impaired in cells bearing GFP-GFAP R239C, despite the increased recruitment of galectins to damaged lysosomes. Interestingly, evidence from diverse experimental systems indicates that GFAP and/or vimentin may interact with several components of the lysosomal repair machinery, including several galectins, the ESCRT component Alix and the galectin-interacting autophagy receptor TRIM16 [47, 67–70]. Nevertheless, further research is needed to elucidate which stages of these processes may be defective.

In summary, our work shows that expression of the AxD GFAP R239C mutation severely affects lysosomal morphology, distribution and function, as well as the ability of these organelles to withstand damage. Although the exact pathway leading to these alterations is still unexplored, our data suggest that lysosomal malfunction may be an important factor to consider in AxD cellular pathology, as it could contribute to failure of autophagy and proteostasis defects, which are key features of this disease. Moreover, together with previous evidence, our results show that not only basic cellular functions appear affected by the presence of the AxD GFAP mutant, but the ability of the cell to cope with various kinds of stress [18, 71]. This recalls the association observed in the clinic between situations provoking stress, such as infections, fever or head trauma and the onset or exacerbation of AxD symptoms (reviewed in [71]), which could inspire strategies for avoiding potentially aggravating factors. Furthermore, a deeper knowledge of AxD cellular pathology arising from several experimental models will hopefully contribute to the design and evaluation of therapeutic approaches.

## Abbreviations

AxD: Alexander disease
GFAP: glial fibrillary acidic protein
LTR: LysoTracker Red
LAMP: lysosomal associated membrane protein.

## Acknowledgements

We are indebted to the personnel from the Laser Confocal and Multidimensional in vivo Microscopy facility at Centro de Investigaciones Biológicas Margarita Salas for expert assistance with confocal microscopy, as well to the personnel of the Flow Cytometry facility for performance of cell sorting. We acknowledge the valuable technical assistance of Paula Martínez-Cenalmor. We are grateful to the Astromad (Fundación “la Caixa”, LCF/PR/HR21/52410002) and Alexander (EJPRD2019 COFUND-EJP N° 825575) consortia for fruitful interactions and discussion. We acknowledge feedback from the COST Action EpiLipidNet CA19105.

## Funding

This work was supported by Fundación “la Caixa”, grant “Astromad”, ref. LCF/PR/HR21/52410002, and in part by Grants PID2021-126827OB-I00 and RTI2018-097624-B-I00 from MCIN, Spain, funded by MCIN/AEI/10.13039/501100011033 and by “ERDF A way of making Europe”. AVP was supported by a contract with reference BES-2016-076965 granted by MCIN/AEI 10.13039/501100011033 and ESF “Investing in your future”. NGI is the recipient of a predoctoral contract funded by Comunidad Autónoma de Madrid, Spain, ref. PIPF-2022/SAL-GL-25771.

## Notes

### Competing Interest Statement

The authors have declared no competing interest.

