## Supplementary material for "Lysosomal function, resistance to stress and repair are compromised by expression of the Alexander disease GFAP R239C mutant"

Department of Molecular and Cellular Biosciences. Centro de Investigaciones Biológicas Margarita Salas, CSIC, Spain

Running Title: Lysosomal damage in AxD

**Supplementary figures**

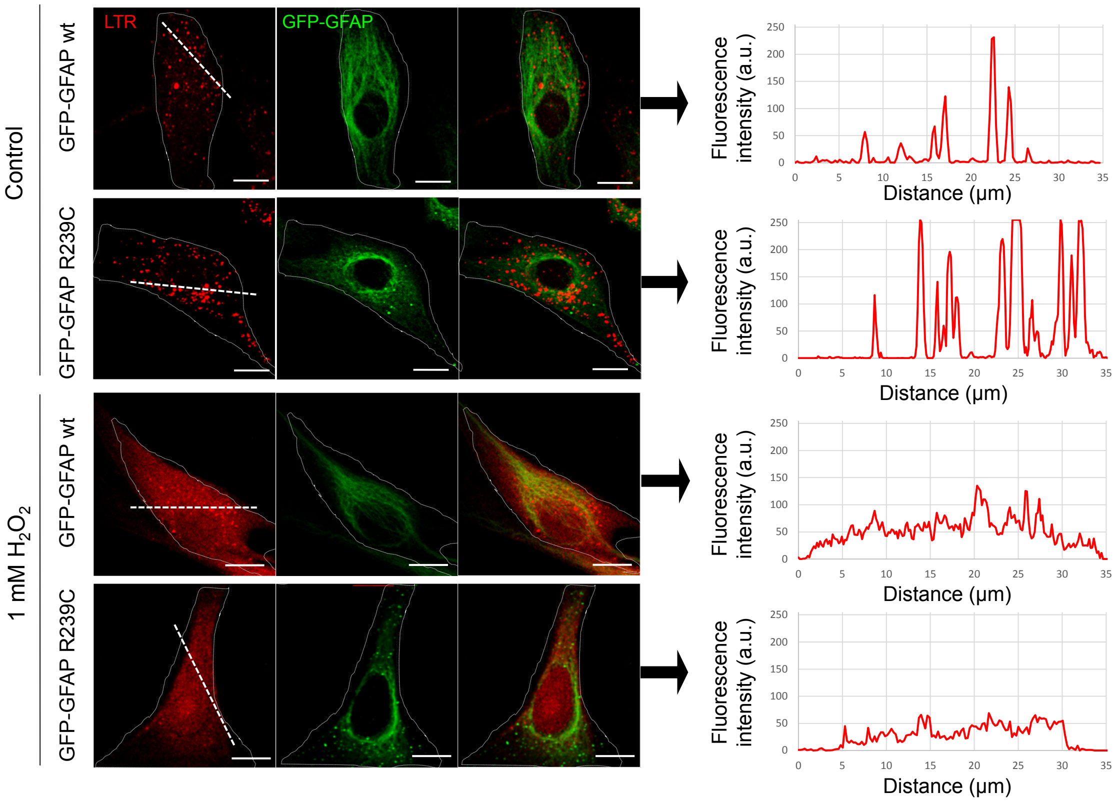


**Suppl. Fig. 1. Effect of $H_2O_2$ treatment on the distribution of lysosomes and lysosomal content followed by LTR staining.** U-87 MG cells expressing GFP-GFAP wt or R239C were incubated in the absence or presence of 1 mM $H_2O_2$ for 30 min, as indicated. LTR staining and GFP-GFAP distribution were assessed by confocal microscopy, and images show individual and merged channels of overall projections. Right panels depict fluorescence intensity profiles of LTR staining along the dotted lines, illustrating the defined pattern of LTR-positive particles in control cells, and the loss of intensity and definition in $H_2O_2$-treated cells, along with a detectable diffuse background, indicative of lysosomal leakage.

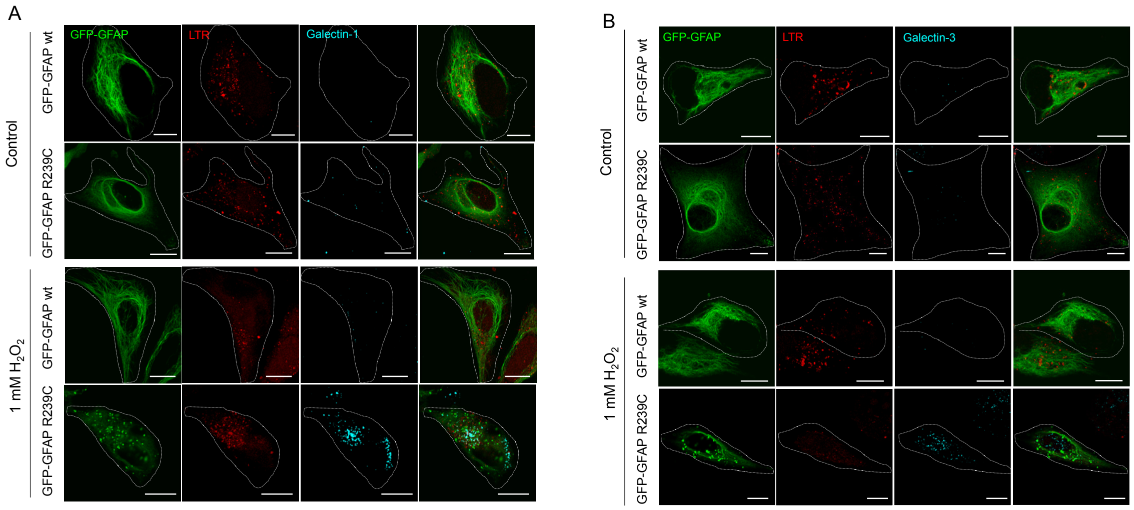


**Suppl. Fig. 2. Detection of galectin-1 and -3 in damaged lysosomes.** Images in panels A and B show the single and merged channels from the experimental conditions described in Fig. 5D and F.

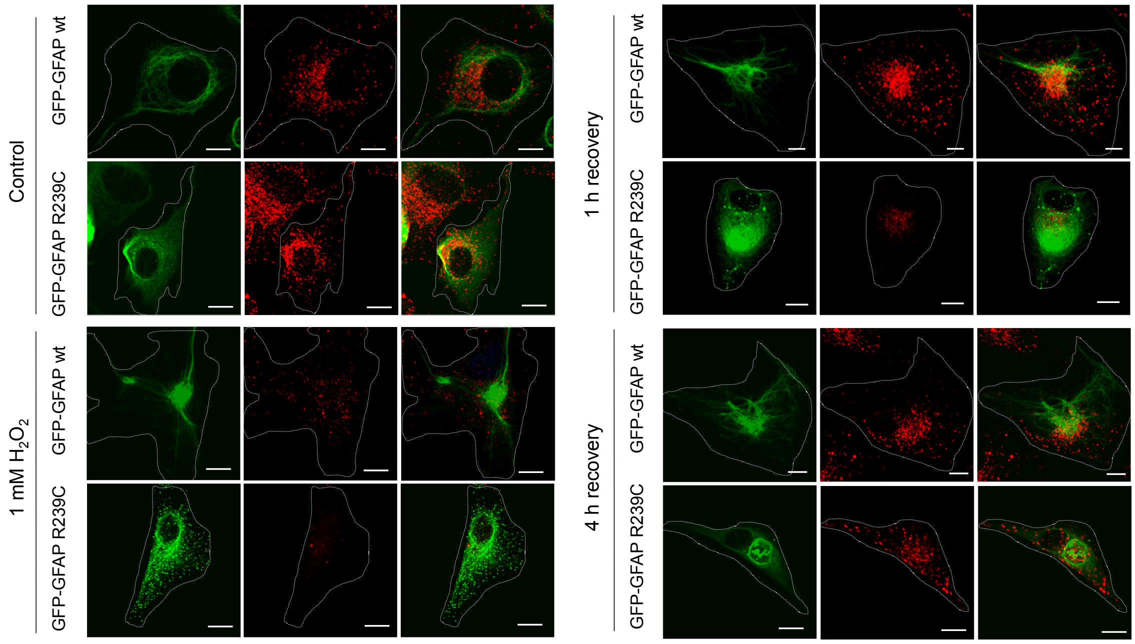


**Suppl. Fig. 3. Lysosomal recovery after oxidative damage.** Images show the single and merged channels from the experiment described in Fig. 7.
